# Spatial transcriptomic analysis of childhood ependymoma implicates unresolved wound healing as a driver of tumor progression

**DOI:** 10.1101/2022.03.29.486280

**Authors:** Rui Fu, Gregory A. Norris, Nicholas Willard, Andrea M. Griesinger, Kent A. Riemondy, Vladimir Amani, Enrique Grimaldo, Faith Harris, Todd C. Hankinson, Siddhartha Mitra, Timothy A. Ritzmann, Richard R. Grundy, Nicholas K. Foreman, Andrew M. Donson

## Abstract

Childhood ependymoma (EPN) is a brain tumor that has seen limited improvements in outcome over past decades. The underlying cellular components of EPN have recently been revealed by single cell RNA-sequencing (scRNAseq), providing biological insights. Here we use spatial transcriptomics to comprehensively chart gene expression across the cellular landscape of posterior fossa subgroup A (PFA) EPN, the commonest and most deadly EPN variant, providing novel resolution of cellular heterogeneity and cellular interaction. We reveal that PFA are comprised of epithelial and mesenchymal histological zones each containing a diversity of cellular states. These include co-existing and spatially distinct undifferentiated progenitor-like clusters - a quiescent mesenchymal zone population, and a second highly mitotic progenitor population that is restricted to hypercellular epithelial zones. We show that myeloid cell interaction is the leading cause of mesenchymal transition in PFA, occurring in zones spatially distinct from hypoxia-induced mesenchymal transition, and these distinct EMT-initiating processes were replicated in in-vitro models of PFA. Collectively, our transcriptomic and functional analyses mirror the processes of normal wound healing where PFA mesenchymal and epithelial zones interact with immune subpopulations in cycle of persistent tissue damage response and mitogenic re-epithelialization signals. Spatial transcriptomics advances our understanding of PFA biology, implicating a “wound that will not heal” process as a driver of tumor progression, a new concept that could provide novel targets for effective therapeutic intervention.

**Significance:** Spatial transcriptomic analysis of the ependymoma tumor microenvironment identifies novel cell populations and interactions, implicating unresolved wound healing as a clinically actionable driver of tumor progression in this refractory childhood brain tumor.

## Introduction

Brain tumors are the leading cause of cancer death in childhood (1). Ependymoma (EPN) is the second most common malignant brain tumor in children (9%) and improvements in outcome have stalled. Despite current standard treatments consisting of surgical resection and adjuvant radiotherapy, over 50% of children with EPN will relapse and most of these will ultimately die, following a protracted course of multiple recurrences (2,3). Clinical trials utilizing chemotherapy at relapse have failed, largely due to the lack of biology-driven trials (4). There is an urgent need for innovative and rational therapies based specifically on identification of factors driving EPN tumor progression.

Recent advances in our understanding of childhood EPN have relied upon molecular, genomic, transcriptomic and methylomic techniques. Molecular stratification has delineated two major EPN subgroups defined by anatomic location - supratentorial and posterior fossa – the latter being more common in childhood and largely comprised of posterior fossa subgroup A (PFA). In recent years the cellular heterogeneity of the EPN tumor microenvironment (TME) has been charted using single cell RNA-sequencing (scRNAseq), identifying multiple neoplastic cell types that include undifferentiated progenitors, ependymal- and mesenchymal-differentiated progeny (5,6). In the present study, we connect microscopy, molecular, and single cell approaches by utilizing spatial transcriptomics (ST) to identify novel mechanisms that drive PFA EPN progression.

## Methods

### Tumor acquisition and processing

Surgical material was collected at our institution at the time of surgery with consent (COMIRB 95-500) and snap frozen for ST analysis (n=14) (Supplementary Table 1). Samples were diagnosed as EPN by standard clinical pathology workup and PFA molecular subgroup assignment was performed using a DNA methylome-analysis based classification portal (https://www.molecularneuropathology.org/mnp). Our rationale for samples selection was to provide representative samples from PFA1 and PFA2 classifications, samples with chromosome 1q gain, samples from patients less than 1 year of age and samples from both WHO grade III and WHO grade II pathological grades of EPN. PFA patient sample FFPE sections corresponding to ST samples and PFA patient sample mouse xenografts were obtained from archival material at our institution for IHC analyses.

### 10x Visium sample preparation

Tissue optimization was performed to determine the optimal permeabilization time of 18 min. Samples were sectioned at 10μm on a Cryostar NX70 cryostat (Thermo Fisher Scientific), fixed with methanol, stained with H&E, and imaged on an Evos M7000 with brightfield settings. Following image capture, capture slides were permeabilized and processed to generate RNA libraries following 10x Visium protocol. Libraries were sequenced to a depth of 70,000 read pairs per spot calculated from the image, on the Novaseq6000 (Illumina) sequencer with 151×151 bp runs.

### Spatial transcriptomics data analysis

Sequencing data were processed with Space Ranger (10x genomics, v1.2.1), followed by further analysis in R using the Seurat (v4.0.1) tool suite. Spots were filtered to ensure the number of genes detected between 50 and 15000, and less than 50% of UMIs mapped to mitochondrial genes. After initial SCTransform normalization on each sample and principal component analysis on merged data, sample integration was performed with Harmony (v1.0) using 30 principal components and theta=2. UMAP dimension reduction and shared nearest neighbor clustering were carried out on 30 principal components, and clustering results at different resolution settings were explored through Clustree (v0.4.3) visualizations.

Cell type identity of clusters was defined in 3 ways: manual inspection of key markers, automated inference by using previously published scRNA-seq transcriptomes (5) as reference using the R package clustifyr (v1.7.1), and Jaccard index calculation of marker gene overlap of clusters and previously reported markers. Differential gene expression was defined by Wilcoxon test, with thresholds of adjusted p value </= 0.01 and log2 fold change of over 0.25 (Supplementary data 1). DAVID (Database for Annotation, Visualization, and Integrated Discovery: https://david.ncifcrf.gov/; version 6.8) was used to measure enrichment of GOterm Direct genesets in subpopulation signatures, providing gene ontology mappings directly annotated by the source database. Cell type identity overlays on H&E image were produced in Loupe Browser (10x Genomics, v5.1.0). Distance calculations between spots and clusters were performed with clustifyr function calc_distance. Cell cycle phases were inferred by Seurat function CellCycleScoring.

### Immunohistochemistry

Immunohistochemistry (IHC) was performed on 5-µm formalin-fixed, paraffin-embedded tumor tissue sections using a Ventana autostainer. IHC for COL9A2 (Thermo PA5-63286, 1:50), Ki-67 (DAKO MIB-1, 1:400), Iba1 (Waco 019-19741, 1:2000), CA9 (Novus NB100417, 1:850) and VIPR2 (Thermo PF3-114, 1:2000) was performed as previously reported by our laboratory (5,7,8). All immunostained sections were counterstained with hematoxylin. Neuropathological review of staining was then performed (N.W.).

### Perturbation studies of PFA EPN *in vitro*

CD14+ myeloid cells were co-cultured with PFA EPN cell lines MAF-811 and MAF-928(9) *in vitro* for 3 days, followed by measurement of transcriptomic changes in tumor cells. CD14+ monocytes were isolated from normal donor peripheral blood by magnetic bead isolation (EasySep, Stem Cell Technologies) then co-cultured with PFA EPN cell lines MAF-811 and MAF-928. Co-cultured CD14+ monocytes and EPN cell lines were separated using cell culture inserts with 0.4mm pores, that allow for free diffusion of soluble factors (Corning) and prevention of cell type cross-contamination and incubated at a ratio of ∼1:1 for 3 days. RNA was then harvested from EPN cell lines using an RNeasy mini kit (Qiagen). Illumina Novaseq6000 libraries were prepared and sequenced by the Genomics and Microarray Core facility at the University of Colorado Anschutz Medical Campus. High quality base calls at Q30>80% were obtain with 20 million paired-end reads. Sequenced 150bp paired-end reads were mapped to the human genome (CRCh38) by STAR 2.4.0.1. Gene expression, as fragments per kilobase per million (FPKM), was derived by Cufflinks. In addition, RNAseq expression FPKM data from EPN PFA cell lines MAF811 and MAF928 that had been cultured in normoxia or hypoxia (2% O^2^) for 3 days was obtained from a prior study by our laboratory (5). Genes upregulated (fold change >1.25) in treated cells compared to controls for each experiment were identified, and commonly upregulated genes between technical replicates were then subject to hypergeometric analysis to quantify enrichment of the top 50 ST cluster marker genes (Supplementary data 1).

### Statistical analyses

Statistical analyses were performed using R bioinformatics, Prism (GraphPad), and Excel (Microsoft) software. For all tests, statistical significance was defined as P < 0.05.

### Data availability

Visium ST data has been deposited in the National Center for Biotechnology Information Gene Expression Omnibus (GEO) database and are publicly accessible through GEO accession number GSE195661 (https://www.ncbi.nlm.nih.gov/geo/query/acc.cgi?acc=GSE195661). PFA ST data including H&E histology, clusters and gene expression for all 14 samples is available as a browsable internet resource at https://raysinensis.shinyapps.io/spatialshiny/.

## Results

### Spatial transcriptomics reveals that the PFA TME is comprised of epithelial and mesenchymal zones

ST was used to map transcriptomic data onto tumor cellular architecture in PFA using the Visium platform (10x Genomics). We analyzed 14 (11 primary; 3 matched recurrence) snap frozen surgical sections, representing a range of PFA molecular and clinical variables (PFA1/PFA2 subgroup classification, chromosome 1q gain, samples from patients less than 1 year of age and samples from WHO grade II and III) (Supplementary Table 1). ST sequencing data was processed and filtered (Supplementary Fig. 1) resulting in 33,082 spots across 14 samples (∼2,400 spots per sample). Spots from each sample were clustered with batch correction/alignment which identified 23 ST clusters, 18 of which were present in all 14 samples. Morphologically, PFA EPN are typified by areas of varying cell density, occasionally arranged in rosettes (Fig. 1A). The histological arrangement of ST clusters mirrors this morphological heterogeneity across the cellular landscape of PFA (Fig. 1B).

**Figure 1.**
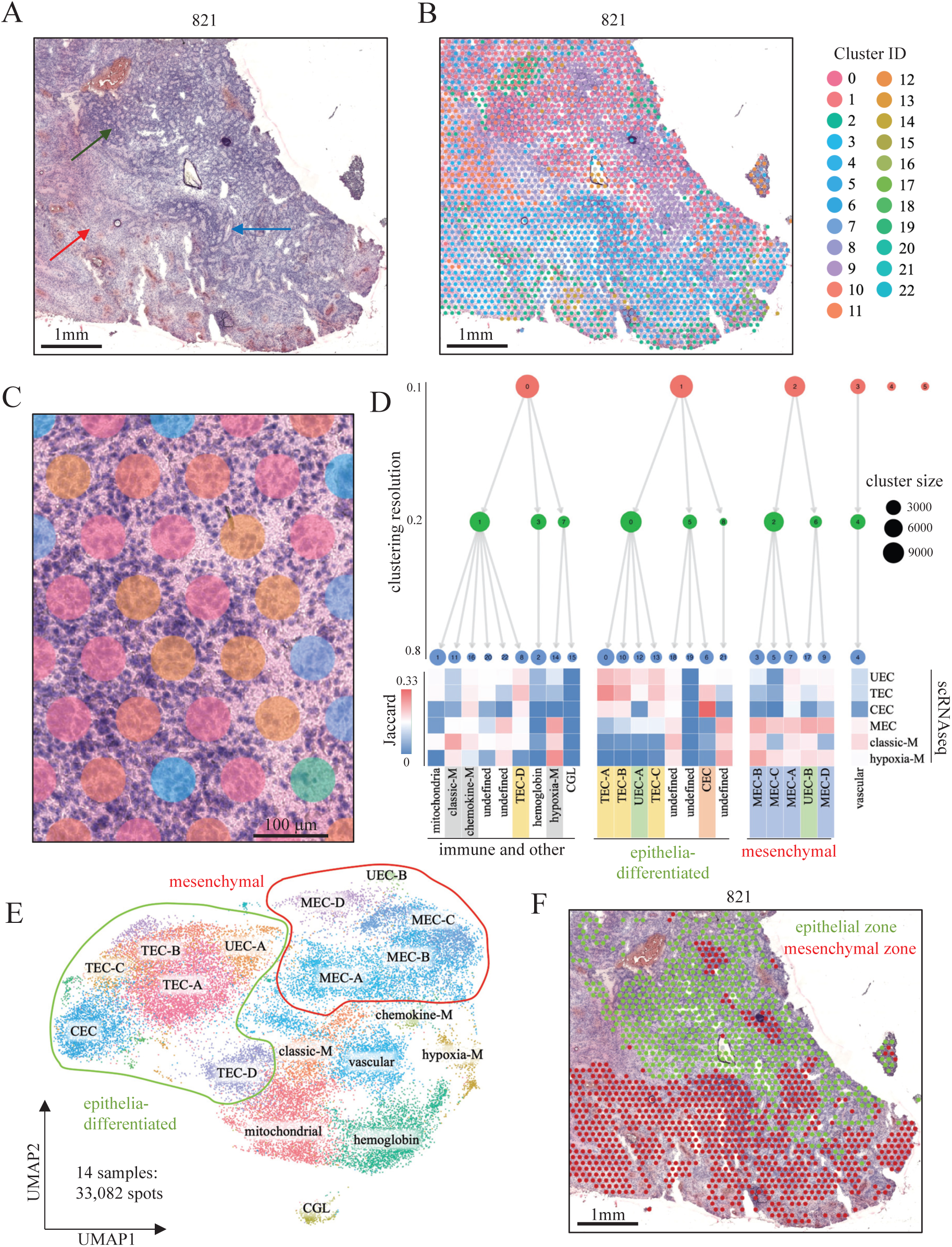
The PFA TME is comprised of major epithelial and mesenchymal transcriptomic zones. **A**, Representative low magnification image of PFA EPN (H&E) in sample 821 showing classic EPN histology, including regions of hypercellularity with prominent perivascular pseudorosettes (blue arrow), areas of increased true ependymal rosettes and epithelial differentiation (green arrow), and paucicellular regions with a lesser degree of differentiation (red arrow). **B**, Overlayed ST clusters (Seurat/Harmony) revealing heterogenous transcriptomic signatures across the TME. **C**, Enlarged magnification highlighting varying numbers of underlying cells per spot (range ∼5 to ∼30). **D**, Sequencing data from 33,082 spots in 14 PFA samples aggregated and processed to identify 23 conserved spot clusters using Seurat with Harmony alignment. Cell type identification of clusters included Jaccard index analysis to calculate marker gene overlap with previously published scRNAseq subpopulation markers, displayed as a heatmap, with clusters annotated accordingly. Clusters were visualized at different resolutions using Clustree analysis which identified 4 major neoplastic and non-neoplastic cluster groups: (i) epithelial-differentiated, (ii) mesenchymal, (iii) immune and other, and (iv) vascular endothelium (abbr: vascular). **E**, UMAP clustering of spatial transcriptomic data demonstrates that neoplastic clusters are arranged into epithelial or mesenchymal groups. **F**, Spatial analysis demonstrates that major neoplastic clusters are commonly arranged into cohesive epithelial and mesenchymal zones that are associated with hypercellular differentiated and paucicellular less differentiated regions respectively. Abbr: CEC, ciliated EPN cells; TEC, transportive EPN cells; UEC, undifferentiated EPN cells; MEC, mesenchymal EPN cells; CGL, cerebellar granular layer.

Transcriptomic profiles from each spot cluster were examined using 3 methods to characterize the identities of cells that underlie distinct clusters: (i) Jaccard index comparison of ST cluster marker genes with previously identified PFA scRNAseq subpopulation marker genes(5); (ii) Clustree analysis(10) of ST cluster similarity; and (iii) ontological analysis of ST marker genes. Spots (diameter 55μm) were comprised of multiple cells, ranging from 1-30 cells depending on the underlying cellularity of the histological region (Fig. 1C). Thus, ST spot clusters largely represent conserved composites of cells rather than individual cell types and are not therefore perfectly comparable with transcriptomic profiles of single cells identified using scRNAseq. Despite this caveat, the aggregated results of the 3 characterization methods allowed us to identify a predominant neoplastic or immune cell type identified by scRNAseq in each major ST cluster. Our previous scRNAseq analysis of PFA identified four major neoplastic subpopulations (5). Two ependymal-differentiated cell types were delineated, ciliated EPN cells (CEC) and transportive EPN cells (TEC), respectively characterized by expression of genes and proteins related to normal ependymal functions such as cilia-associated genes CAPS and FOXJ1, or transporter proteins including AQP4 and VIPR2. The mesenchymal subpopulation (MEC; mesenchymal EPN cells) was shown by immunohistochemistry (IHC) of MEC marker CA9 to border necrotic zones. The progenitor population, termed undifferentiated EPN cell (UEC), was seen to associate with MEC by IHC using UEC marker c-fos. These findings were corroborated in a parallel scRNAseq study (6). These four scRNAseq neoplastic cell subpopulations were differentially enriched in ST clusters (Jaccard index analysis) and ST clusters were annotated accordingly (Fig. 1D,E). EPN arise in the ventricles of the central nervous system and are understood to arise from normal ependymal cells, an epithelial cell type that lines the ventricles. TEC/CEC subpopulations therefore represent epithelial-differentiated EPN cell types. Accordingly, Clustering was dichotomized into major neoplastic epithelial and mesenchymal branches, predominated respectively by TEC/CEC and MEC subpopulations (Fig. 1D). Epithelial and mesenchymal ST clusters were also seen to segregate by UMAP 2D projection of all 14 samples (Fig. 1E). The mesenchymal ST cluster branch represents the process of epithelial-mesenchymal transition (EMT) that has been identified in PFA (5,11,12). Histological examination of ST clusters reveals that most samples are comprised of widespread zones of either epithelial or mesenchymal ST spots (Fig. 1F, Supplementary Fig. 2). As expected, epithelial zones are associated with cell morphologies displaying the rosette-rich and highly cellular areas. Conversely, mesenchymal zones are often moderately or sparsely cellular, and associated with areas of necrosis. ST cluster proportions provide an opportunity to quantify subpopulation proportions more reliably than data from scRNAseq, as ST is a direct measure of relatively unmanipulated tissue whereas scRNAseq involves numerous tissue processing steps that are likely to skew cellular proportions. On average, epithelial zone spots were more abundant than mesenchymal in PFA at presentation (average 1.7:1, n=11), but with a high level of variability (range 3.0:1 to 1:5.6). PFA with a high mesenchymal:epithelial ratio contained large cohesive mesenchymal zones that were often greater than 1mm in the smallest dimension, whereas samples with a higher epithelial:mesenchymal ratio harbored mesenchymal spots that were scattered within epithelial zones (Supplementary Fig. 2). Both epithelial and mesenchymal zones contained multiple ST subclusters but with significant spatial and molecular distinctions, details of which are elaborated in subsequent sections.

The remaining ST clusters were largely associated with non-neoplastic cell identities. A singular and relatively abundant ST cluster was associated with vascular endothelium (VE) gene expression including *VWF* and *PECAM1* (Fig. 1D,E; Supplementary Data 1,2), occupying an exclusive Clustree branch (Fig. 1D). This finding highlights an additional strength of ST, as this cell type was not identified in prior single cell RNAseq analyses, likely due to the cohesive nature of VE precluding mechanical disaggregation into single cells required for successful capture. A number of ST clusters were predominated by myeloid immune cell gene signatures, distinguished by either classic macrophage/microglia markers such as HLA-class II, hypoxia-associated ontologies, or chemokine expression (Fig. 1D,E; Supplementary Data 1,2) which are elaborated in a subsequent section. Two other ST clusters were predominated either by hemoglobin gene signatures, which were subsequently shown to reside in areas with histological evidence of bleeding (hemosiderin), or by high expression of mitochondrial genes. A single sample (928_2) contained an area of normal brain adjacent to tumor, predominated by cerebellar granular layer (CGL) cells that was seen as a discrete ST cluster (Fig. 1E). This cluster, along with the 6 least abundant ST clusters that were only present in 1 or 2 samples each, were excluded from the remainder of this study. PFA ST data including H&E histology, clusters and gene expression for all 14 samples is available as a browsable internet resource at https://raysinensis.shinyapps.io/spatialshiny/.

### Mesenchymal zones are comprised of spatially restricted EMT stages in PFA

The process of EMT in PFA (11,12) was further elaborated in our recent scRNAseq study, showing that EMT could be induced in PFA cell lines under hypoxic culture conditions (5). However, initiation of EMT *in situ* can potentially be triggered by a number of extrinsic and intrinsic stimuli in addition to hypoxia, including nutrient deprivation, cellular stress, and tissue damage (13). The ST PFA mesenchymal zone is comprised of 5 ST clusters that are enriched for MEC scRNAseq markers (Fig. 1D). Distinctions between mesenchymal zone ST clusters potentially represent different EMT stimuli, or alternatively distinct transition states in the EMT process. To test these hypotheses, we examined differences between mesenchymal zone ST clusters with respect to gene expression (Supplementary Data 1,2) and spatial arrangement (Fig. 2A; Supplementary Fig. 3)

**Figure 2.**
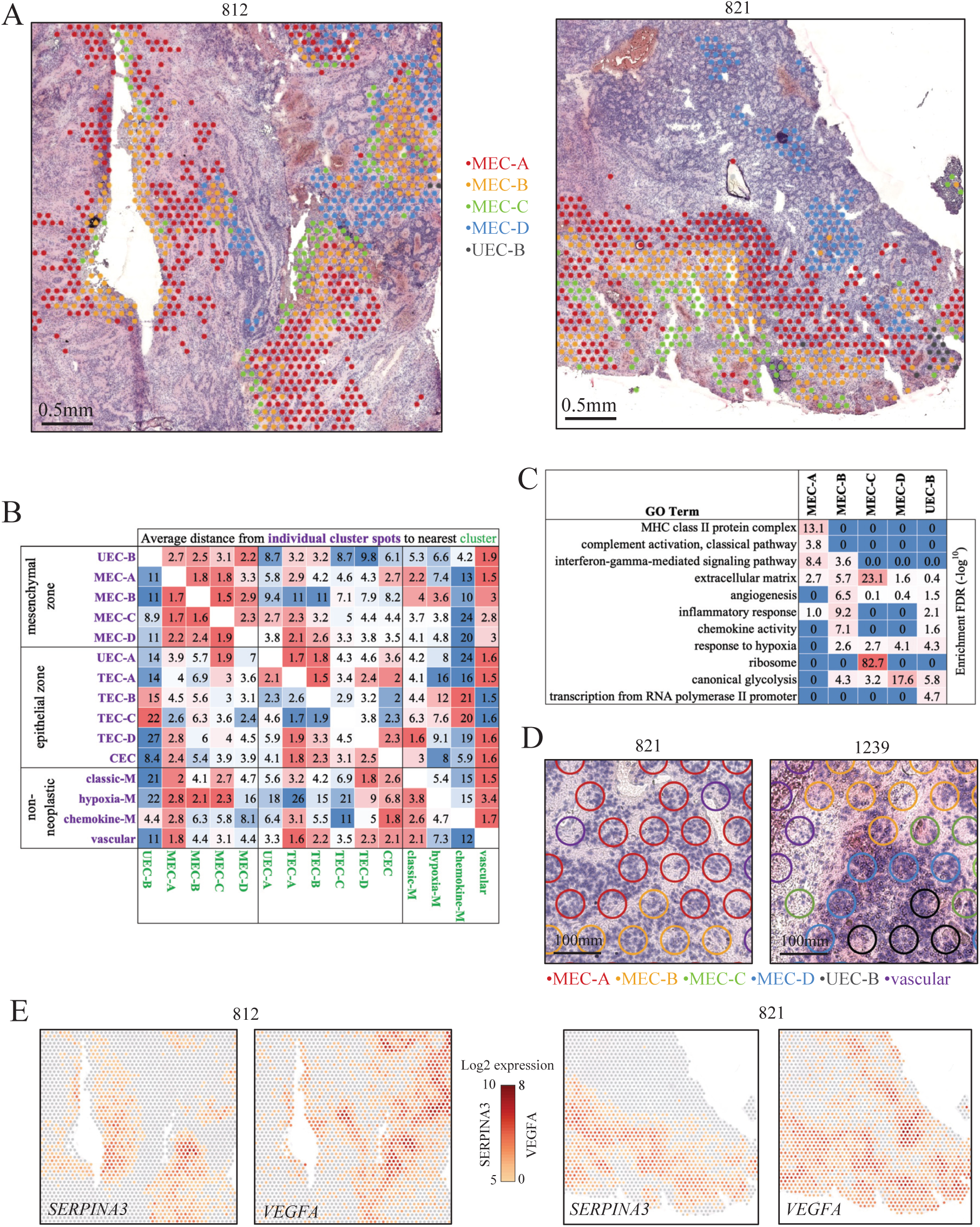
Spatial transcriptomics delineates distinct stages of EMT in PFA. **A**, Representative spatial arrangement of mesenchymal zone clusters in two PFA samples. **B**, Heatmap of average individual cluster spot (y-axis) distances to nearest cluster (x-axis). Distance is measured as number of spots apart (inter-spot distance = 100μm). **C**, Top gene ontology enrichments for mesenchymal zone clusters. **E**, Representative spatial gene expression of key markers of MEC-A (*SERPINA3*) and MEC-D (*VEGFA*). Abbr: CEC, ciliated EPN cells; TEC, transportive EPN cells; UEC, undifferentiated EPN cells; MEC, mesenchymal EPN cells.

Of the 5 mesenchymal zone clusters, one cluster was distinguished by particularly high expression of early response genes such as *FOS* and *JUN* (Supplementary Data 1) that are also markers of the scRNAseq UEC progenitor subpopulation (5), and was thus labelled UEC-B. The remaining 4 mesenchymal zone ST clusters were labelled MEC-A, MEC-B, MEC-C and MEC-D in order of abundance. Our prior scRNAseq study had used c-fos IHC to map the spatial location of UEC in relation to other neoplastic subpopulations, revealing an association of UEC with MEC in perinecrotic and perivascular areas (5). Accordingly, UEC-B were seen to reside in proximity to mesenchymal zone clusters (Fig. 2A; Supplementary Fig. 3). Spatial proximity between cell types is an important factor in their biological roles within the TME. We therefore systematically calculated inter-spot distances across all 18 major ST clusters in all 14 samples. This confirmed the closer spatial relationship of UEC-B with other mesenchymal zone clusters than epithelial zone clusters (Fig. 2B). Ontological analysis revealed that UEC-B is distinguished from other mesenchymal zone subclusters by ontologies related to transcriptional activation of the Pol-II promoter, but shared response to hypoxia and canonical glycolysis with specific MEC subclusters, notably MEC-D (Fig. 2C; Supplementary Data 2), consistent with the commonalities between UEC-B and MEC-D seen by Clustree analysis (Fig. 1C).

The most abundant mesenchymal zone subcluster MEC-A is distinguished by enrichment of immune genes related to MHC-class II and complement (Fig. 2C; Supplementary Data 1,2). MEC-A cluster complement-related marker genes were largely restricted to the neoplastic MEC subpopulation rather than immune cells when mapped to scRNAseq data (Supplementary Fig. 4). These include key complement component 1 genes *C1S, C1R* and *CFB*, suggesting a neoplastic cell-intrinsic activation of the classic complement cascade. Serine protease inhibitor *SERPINA3* is the top marker in MEC-A, playing a potentially cytoprotective role against complement component 1 factors that are serine proteases. Conversely MEC-A MHC class-II genes (*HLA-DRA, HLA-DRB1*) and complement cascade component *C3* marker genes were more restricted to scRNAseq myeloid cells (Supplementary Fig. 4), demonstrating that MEC-A ST subclusters are comprised of a mixture of MEC and myeloid cells, as also seen by Jaccard index analysis (Fig. 1C).

MEC-D cluster spots commonly presented as islands within hypercellular epithelial zones (Fig. 2A; Supplementary Fig. 3) and were microscopically distinguished from other MEC subclusters by more hypercellular histology (Fig. 2D). Glycolytic genes, including *GAPDH, TPI1* and *LDHA*, were enriched in MEC-D (Fig. 2C; Supplementary Data 1,2). Upregulation of glycolytic metabolism is a hallmark of cells responding to hypoxia, and hypoxia-related genes, in particular angiogenic *VEGFA*, are significantly enriched in MEC-D. This suggests that MEC-D arises as a result of hypoxic and/or metabolite depleted conditions that occur in hypercellular, hyperproliferative, epithelial zones. MEC-C is equivalent to MEC-D with respect to enrichment of hypoxia-related genes such as *VEGFA* and *CA9* (Fig. 2C), but in contrast has a predominantly low cell density, and is often seen in necrotic zones (Fig. 2D; Supplementary Fig. 3).

MEC-B is distinguished by genes related to extracellular matrix remodeling, including NF_K_B components (*NFKB1, NFKB2, NFKBIA, NFKBIZ*), cytoprotective protease inhibitors (*SERPINs* and *TIMP1*), chitinases (*CHI3L1, CHI3L2*), metalloproteases (*MMP7, MMP9*) (Supplementary Data 1), and extracellular matrix genes (Fig. 2C; Supplementary Data 1,2). Additionally, MEC-B was distinguished by an enrichment of genes involved with the inflammatory response characterized by chemokines *CXCL1, CXCL8* (*IL8*) and *CCL2* (Fig. 2C; Supplementary Data 2). MEC-B is predominately located in moderately cellular areas with less evidence of residual ependymal-differentiated architecture than MEC-A or MEC-D (Fig. 2D; Supplementary Fig. 3).

To further elucidate the biology of different MEC subclusters, we examined their spatial relationships. MEC subclusters commonly reside in neighboring histological zones (Fig. 2A; Supplementary Fig. 3) with distinct tissue morphology, supporting the hypothesis that the MEC subclusters represent discrete stages in mesenchymal differentiation. MEC cluster zones are often arranged in a conserved order of consecutive waves radiating away from epithelial zones. This striking spatial arrangement suggests a spatial lineage trajectory, potentially representing stages in the EMT with a sequence commonly starting with MEC-A, followed by MEC-B and finally MEC-C (Fig. 2A; Supplementary Fig. 3). Proximity analysis indicates that MEC-D and MEC-A are the most distantly separated MEC subclusters (Fig. 2B), which combined with spatially disparate marker gene expression (Fig. 2E, Supplementary Data 1) suggest that MEC-D and MEC-A represent separate EMT pathways in PFA. Based on gene expression characteristics of tissue remodeling and less epithelial-differentiated, paucicellular and necrotic histology, we hypothesize that MEC-B and MEC-C zones represent later stages in the EMT process.

### Infiltrating myeloid immune cells are spatially related to and interact with specific mesenchymal zone clusters

Three major ST clusters were characterized by immune cell marker gene expression, and these clustered with other non-neoplastic cell types that include vascular endothelium and red blood cells (Fig. 1C). The most abundant immune ST cluster showed significant marker gene overlap with a myeloid scRNAseq subpopulation with gene expression characteristic of classical M1 polarization (Fig. 1C). This included phagocytosis, MHC-class II and complement activation genes (Fig. 3A; Supplementary Data 1,2), and the ST cluster was therefore labelled classic myeloid (classic-M). In contrast, the second most abundant immune cluster overlapped with a myeloid scRNAseq subpopulation with a hypoxia-response gene expression signature (Fig. 1C). Gene ontologies associated with this ST cluster were primarily inflammatory response-related, consisting largely of chemokines involved with neutrophil chemotaxis, and was labeled hypoxia myeloid (hypoxia-M). The smallest myeloid ST cluster, distinguished by particularly high expression of monocyte chemoattractants *CCL3* and *CCL4* (Supplementary Data 1), was labelled chemokine myeloid (chemokine-M).

**Figure 3.**
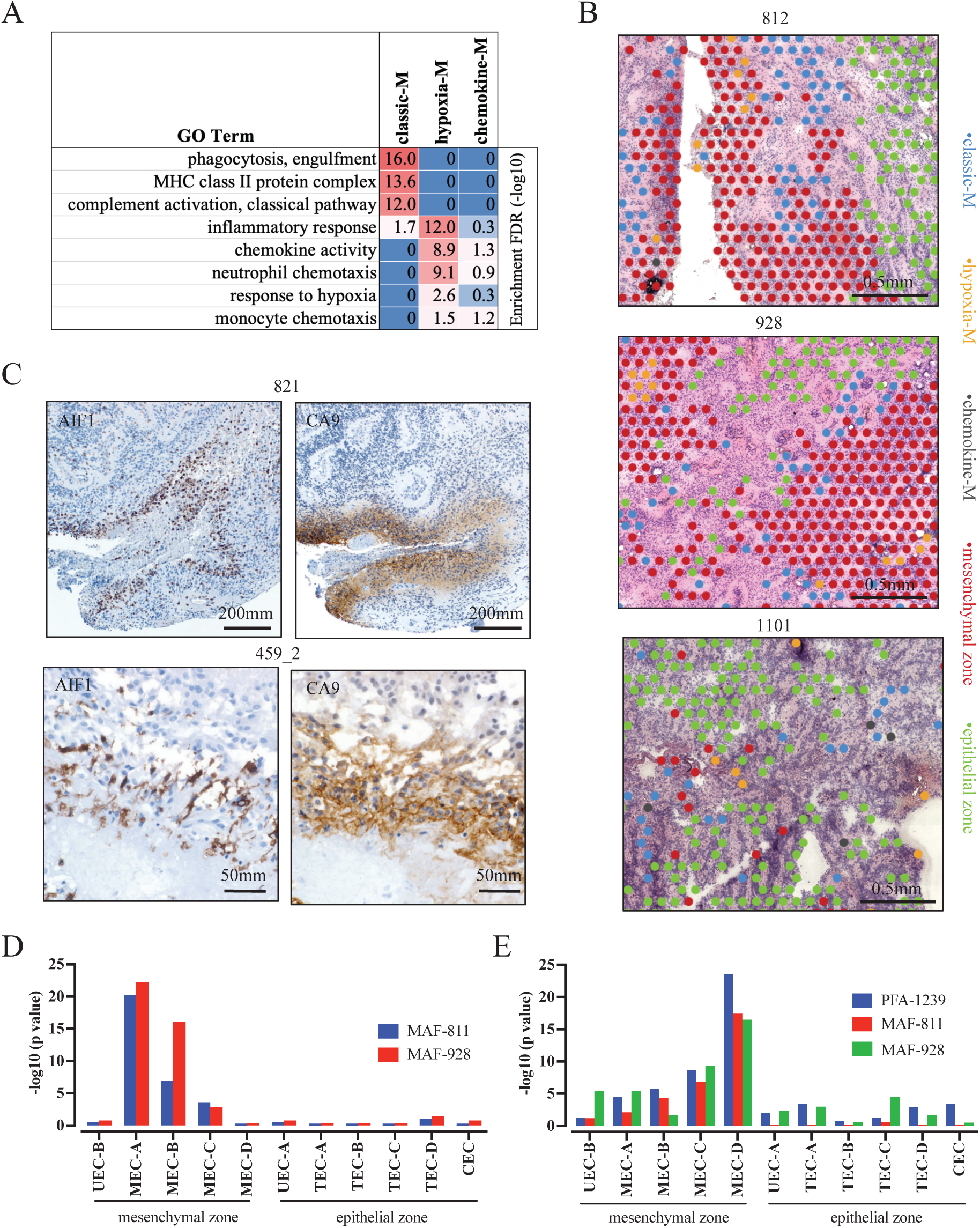
Myeloid lineage immune cells are spatially related to and interact with specific mesenchymal ST clusters in PFA. **A**, Top gene ontology enrichments for mesenchymal zone clusters (hypergeometric test FDR (–log10)). **B**, Representative spatial arrangement of immune clusters with respect to mesenchymal and epithelial zone clusters in three PFA samples. **C**, Representative IHC in sequential sections showing adjacent localization of expression (brown) for markers of classic-M (AIF1) and mesenchymal zones (CA9) in two PFA samples. **D**, Upregulation of ST cluster marker genes in PFA cell lines MAF-811 and MAF-928 after exposure to CD14+ myeloid cells using a modified co-culture system where tumor and immune cells are separated by semi-permeable membranes that allow free movement of soluble factors. After 3 days RNA was harvested and sequenced (RNAseq) to identify upregulated genes compared to untreated cell lines. Commonly upregulated genes (fold change > 1.25) between replicates for each perturbation were then compared to mesenchymal and epithelial zone ST cluster markers (top 50) to identify cluster specific marker gene enrichment (hypergeometric analysis p-value (-log10)). **E**, Upregulation of ST cluster specific genes in PFA cell lines (MAF-811 and MAF-928) and short term PFA culture (PFA-1239) after propagation in 2% hypoxia for 3 days was obtained from a previous study (5) and analyzed as above. Abbr: CEC, ciliated EPN cells; TEC, transportive EPN cells; UEC, undifferentiated EPN cells; MEC, mesenchymal EPN cells.

Spatial analysis revealed different patterns of classic-M and hypoxia-M spot localization in relation to mesenchymal zones. We observed cohesive bands of classic-M ST cluster spots bordering large mesenchymal zones (Fig. 3B; Supplementary Fig. 5). This distinctive cellular arrangement was confirmed using IHC with classic-M marker AIF1 (Iba1) and MEC marker CA9, revealing dense bands of myeloid and MEC cells lining areas of necrosis (Fig. 3C). Proximity analysis of immune and neoplastic ST clusters revealed that classic-M was most closely associated with MEC-A (Fig. 2B). Hypoxia-M are also associated with mesenchymal zones, but in contrast to classic-M are embedded within mesenchymal areas rather than at their borders (Supplementary Fig. 5) suggesting a potential role in later stages of EMT, such clearance of cellular debris.

The spatial colocalization of classic-M with cells at the border of MEC-A suggests a potential immune role in initiation of EMT. We performed a functional test of this hypothesis by co-culturing CD14+ myeloid cells with PFA EPN cell lines MAF-811 and MAF-928(9) *in vitro* for 3 days, followed by measurement of transcriptomic changes by RNAseq. Cell lines co-cultured with CD14+ myeloid cells showed significant upregulation of mesenchymal zone cluster signature genes but not epithelial zone clusters (Fig. 3D). Specifically, we observed a predominant myeloid-mediated upregulation of MEC-A marker genes (including *HLA-DRA, C1S, C1R, SERPINA3, CHI3L1*) in both cell lines, whereas no significant enrichment of MEC-D signature genes was observed. In contrast, propagation of PFA cell lines (MAF-811 and MAF-928) and a short term culture (PFA-1239) in hypoxic conditions (3 days at 2% O^2^), previously shown to upregulate mesenchymal genes (5), specifically upregulated MEC-D marker genes (including CA9, LDHA) (Fig. 3E). These results further support the co-existence of two distinct EMT pathways in PFA, where MEC-A is initiated by myeloid cell interaction, potentially caused by disruption of normal epithelial cell architecture, whereas MEC-D arises under hypoxic conditions that arise in the context of rapid tumor proliferation. Hypoxia was previously considered to be the major cause of EMT in EPN (5), but the present study reveals that at tumor presentation myeloid cell-associated EMT is significantly more widespread, with MEC-A constituting on average 2.6% of tumor volume versus 1.1% for MEC-D (p=0.016, paired 2-tailed t-test, n=11).

### Identification of a novel proliferative progenitor cluster in PFA epithelial zones

ST epithelial zones are comprised of 6 major ST clusters that are present in all 14 PFA samples analyzed. These includes 3 ST clusters with similarity to the scRNAseq TEC subpopulation, labeled TEC-A, TEC-B and TEC-C in order of abundance (Fig. 1D). An isolated TEC ST cluster was located with Clustree branch largely comprised of non-neoplastic ST clusters despite a high enrichment of TEC scRNAseq marker genes and was labeled TEC-D (Fig. 1D). Unexpectedly, one epithelial zone ST cluster expressed genes restricted to the UEC scRNAseq subpopulation, specifically *ELN, MFAP2, MDK* and *COL9A2* (Supplementary Fig. 6; Supplementary Data 1), ontologically related to elastic fiber (Supplementary Data 2). This cluster was more abundant than the mesenchymal zone progenitor subpopulation UEC-B and accordingly, this cluster was named UEC-A. UEC-A harbored a distinct gene expression profile from UEC-B (Fig. 4A) and was distinguished from other epithelial zone clusters by enrichment of ribosome gene expression (FC=60.1, p<0.0001) (Supplementary Data 1), indicating a relatively high level of protein synthesis. Proximity analysis also demonstrated that UEC-A and UEC-B were spatially distant, with UEC-A intermixing with epithelial zone clusters (Fig. 2B; Fig. 4B,C; Supplementary Fig. 7) unlike UEC-B that were restricted to mesenchymal zones (Fig. 2A,B; Supplementary Fig. 3). UEC-A spots were often associated with differentiated epithelial features such as true ependymal rosettes (Fig. 4B,C).

**Figure 4.**
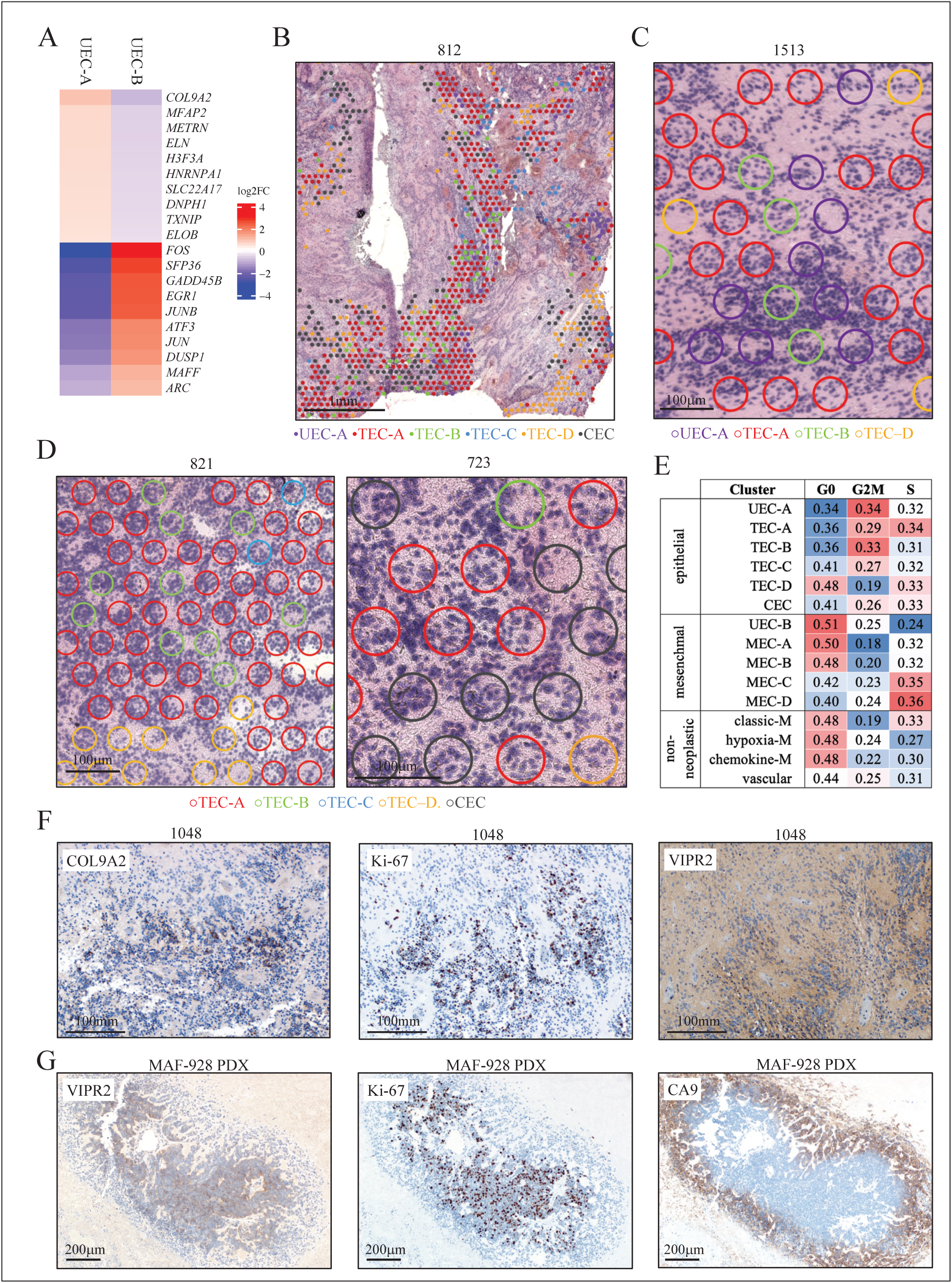
PFA epithelial zones are constituted by progenitor-like and ependyma-differentiated subclusters. **A**, Heatmap of top differentially expressed genes between UEC-A and UEC-B clusters. **B**, Low magnification showing typical spatial distribution of epithelial zone clusters. Representative histology illustrating (**C**) hypercellular histology of UEC-A, and (**D**) TEC and CEC clusters. **E**, Cell cycle phase proportions in neoplastic and non-neoplastic clusters. **F**, Representative IHC images of UEC-A marker COL9A2, mitosis marker Ki-67 and TEC marker VIPR2 in sequential sections. **G**, Low magnification histology of VIPR2, Ki-67 and MEC marker CA9 in sequential sections. Abbr: CEC, ciliated EPN cells; TEC, transportive EPN cells; UEC, undifferentiated EPN cells; MEC, mesenchymal EPN cells.

A single ST cluster (labeled CEC) corresponding to the scRNAseq CEC subpopulation was accordingly enriched for genes related to cilia formation (Supplementary Data 1,2) and often associated with ependymal rosettes (Fig. 4D). Spatially, CEC tend to localize within epithelial zone adjacent to mesenchymal zones (Supplementary Fig. 7).

The predominant epithelial zone ST cluster TEC-A is the most abundant cluster identified in all PFA samples, and similar to UEC-A were commonly associated with PFA differentiated histological areas (Fig. 4B,C; Supplementary Fig. 7). *HES6*, a nervous system development gene, is the top marker gene in TEC-A, along with a number of genes previously identified in TEC scRNAseq analysis including transporters *VIPR2* and *AQP4* (Supplementary Data 1). Individual TEC-B ST cluster spots were intermixed with TEC-A in differentiated areas (Fig. 4B-D; Supplementary Fig. 7) but distinguished from TEC-A by expression of *CTNNB1, DDX3X* and *LRCC75A* (Supplementary Data 1). TEC-D was generally distinguished from other TEC subclusters by low cellularity (Fig. 4B-D) and a relatively high expression of *GFAP* (Supplementary Data 1).

Analysis of cell cycle phase gene expression revealed that epithelial zone ST clusters were overall more proliferative than mesenchymal zone or non-neoplastic clusters (Fig. 4E). The most proliferative of all ST clusters was UEC-A, concordant with the predicted high protein synthesis inferred by ribosome gene expression. In contrast, mesenchymal associated UEC-B had the highest proportion of cells in G0 cell cycle phase, suggesting a quiescent phenotype. We confirmed the high proliferative rate in UEC-A using IHC for UEC-A marker COL9A and TEC-A marker VIPR2 in relation to mitosis marker Ki-67, showing co-localization of COL9A2 and Ki-67 in hypercellular regions (Fig. 4F). IHC also confirmed the relatively high proliferative rate of epithelial zones versus to CA9+ mesenchymal zones (Fig. 4G).

## Discussion

Identification and characterization of mesenchymal and epithelial subpopulations using spatial transcriptomics (ST) integrated with scRNAseq allows us to more accurately visualize and chart stages of tumor progression in PFA. This is illustrated by our discovery of co-existing progenitor clusters with distinct phenotypes spatially restricted to either epithelial or mesenchymal TME niches. The phenomenon of coexisting cancer stem cells (CSCs) within the same primary tumor has been identified in human breast cancer (14). This study charted proliferative epithelial-like CSCs and quiescent mesenchymal-like CSCs, showing remarkable molecular and spatial similarity to the progenitor phenotypes in PFA in the present study. We contend that dual PFA progenitor populations are integrally involved in cellular plasticity via EMT. Identification of UEC-A as a highly proliferative progenitor provides a novel target for therapeutic development in PFA.

The increased resolution afforded by ST also revealed the existence of two independent EMT pathways within the TME – one associated with myeloid cell interaction, and a second induced by hypoxia. EMT stages in cancer have previously been characterized using transgenic mouse models, combined with lineage tracing, flow cytometry, multicolor immunofluorescence and scRNAseq (15). Here we use ST to provide an orthogonal approach for delineation of EMT stages *in situ* in autochthonous tumors. Further work will be needed to determine whether the mesenchymal cells as a whole or distinct EMT trajectories contribute to the invasive phenotype of EPN. This is a critical clinically relevant question given the high recurrence rates in EPN attributed to microscopic disease that remains after gross total resection and radiation therapy, and that is potentially related to the invasive aspect of EMT. This hypothesized EMT-associated invasiveness may explain not only recurrence, but also the dire clinical outcomes experienced by children with recurrent EPN in whom standard treatment approaches usually fail (3).

Myeloid-associated EMT in PFA is an inflammatory process that may be triggered by homeostatic perturbations such as breakdown in architecture (loss of structure being an inflammatory event for epithelial tissues) or hypoxia, or alternatively an antitumor response driven by recognition of tumor associated antigens. An antitumor immune process is unlikely, given that PFA ependymoma has a mutational burden below the threshold considered necessary to trigger a robust tumor antigen-driven antitumor immune response. In support of a homeostatic role, the accumulation of myeloid cells around mesenchymal zones in PFA closely resembles myeloid corralling that has been identified in spinal cord injury (16), which forms a barrier to protect surrounding “healthy” tissue from tissue that is undergoing inflammatory homeostatic remodeling.

Thus, in PFA the host immune system may regard the tumor as self, initiating a potentially tumorigenic repair process. Indeed, the collective spatial cellular and molecular insights revealed by ST suggest that PFA ependymoma is a “wound that will not heal” (Fig. 5). The similarities between wound healing and tumorigenesis have been proposed previously (17) but have been difficult to prove. The wound healing process consists of 3 main phases: (i) an inflammatory phase, (ii) a regenerative phase of re-epithelialization, and (iii) a final resolution phase. Our present study has identified PFA processes that mirror the initial inflammatory phase and elements of tissue remodeling inherent in the regenerative stage. The regenerative phase of wound healing also encompasses re-epithelialization, which is achieved by the activity of spatially separated migratory and proliferative epithelial zones. In normal skin, re-epithelialization involves the undamaged proximal epithelial layer, comprised of more differentiated epithelial cells, advancing *en masse* to repopulate the damaged area. This process is driven by a distal proliferative pool of epithelial cells repopulated by an epithelial stem cell. In PFA, we hypothesize that UEC-A mirrors this distal proliferative epithelial progenitor subpopulation. Our data suggest an interactive cycle between epithelial and mesenchymal zones, whereby slowly cycling mesenchymal cells, that arise from epithelial cells as a result of a variety cellular stresses, send signals that re-epithelialize the tumor through activation and proliferation of the epithelial cell proliferative pool. This drives more differentiated epithelial zone cells to “re-epithelialize” the tumor, potentially indicated by the frequent clustering of CEC adjacent to mesenchymal zones. As the tumor never achieves the final resolution phase of wound healing, the cycle of inflammation and re-epithelialization persists resulting in unchecked tumor progression. Investigations testing hypothetical EMT-driven re-epithelialization in PFA may reveal novel therapeutic interventions.

**Figure 5.**
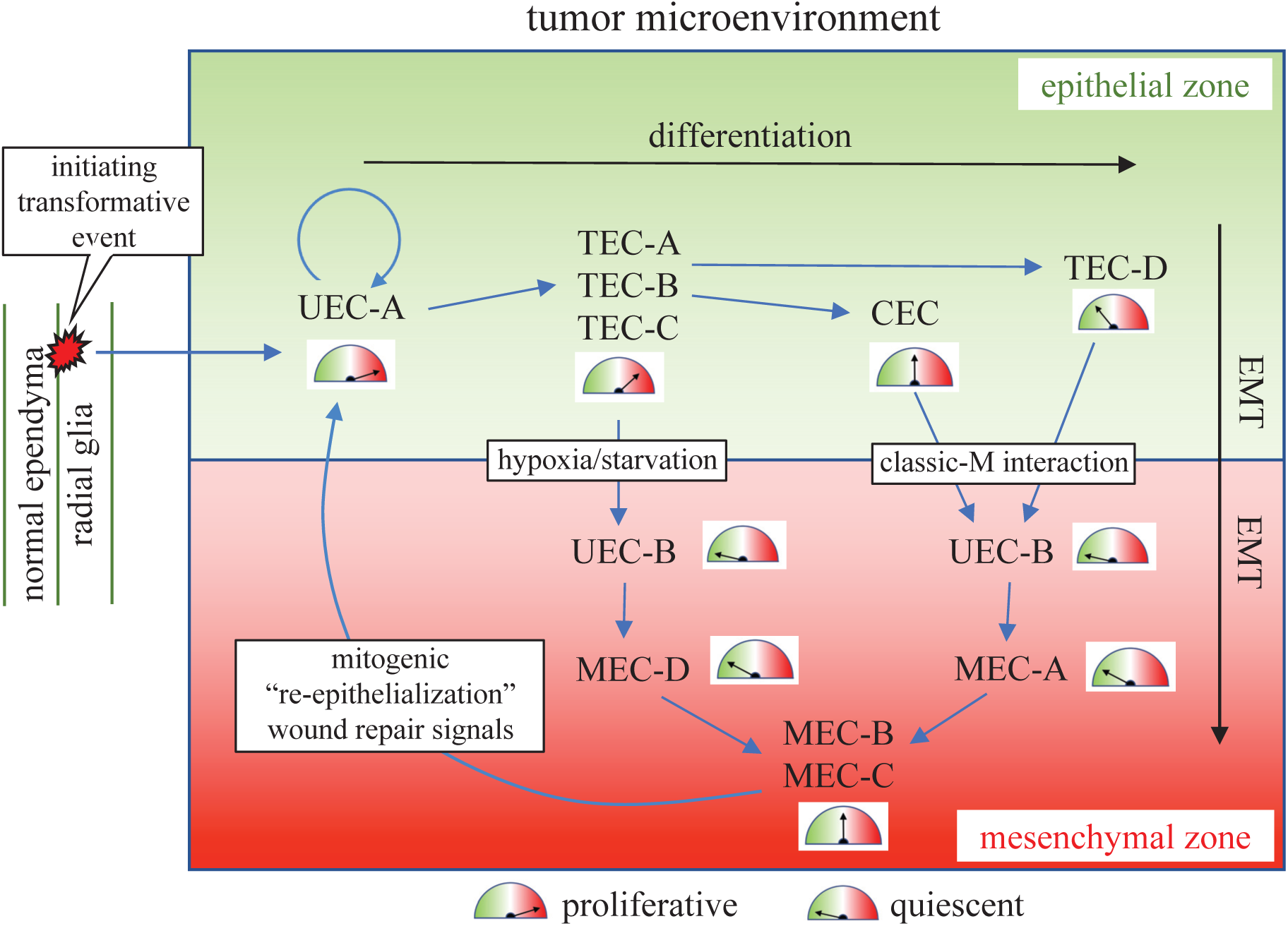
Schematic of PFA neoplastic and immune subpopulation transitions and interactions that drive a cycle of persistent cellular stress and unresolved repair. Spatial cellular and molecular insights revealed by ST suggest that a “wound that will not heal” process is driving tumor progression in PFA. In this model, EPN tumor growth is driven by a highly proliferative epithelial zone progenitor (UEC-A). Cellular stress in epithelial zone cells drives EMT that is mediated by myeloid cell interaction or hypoxic response to generate mesenchymal-differentiated cells (MEC) via a quiescent mesenchymal progenitor (UEC-B). MEC potentially hijack normal wound healing re-epithelialization signals to drive further proliferation in epithelial zone progenitors, resulting in a persistent cycle that drives PFA tumor progression. Abbr: CEC, ciliated EPN cells; EMT, epithelial/mesenchymal transition; MEC, mesenchymal EPN cells; TEC, transportive EPN cells; UEC, undifferentiated EPN cells.

The concept of a “wound that will not heal” in PFA is may also apply to histologically related supratentorial EPNs which are predominantly caused by a *ZFTA*-*RELA* gene fusion (18). This fusion results in constitutive activation of RELA, a key component of the NF_K_B pathway, a critical controller of inflammation and tissue remodeling programs. Thus, persistently dysregulated inflammation and tissue remodeling-driven by the *ZFTA*-*RELA* fusion in supratentorial EPN represents a phenocopy of PFA, suggesting that a common process of unresolved wound healing drives progression in the majority of EPN tumors.

## Acknowledgements

R.F and K.A.R. are supported as informatics fellows of the RNA Bioscience Initiative, University of Colorado School of Medicine. This study was supported by the Tanner Seebaum Foundation, the Morgan Adams Foundation, and the NIH (R01CA237608). The University of Colorado Denver Genomics and Microarray, and Histology Shared Resources are supported by the University of Colorado NIH/NCI Cancer Center (P30CA046934).

## Author Contributions

A.M.D. performed experiments with assistance from A.M.G., V.A., E.G., F.H., S.M. and T.C.H. R.F performed bioinformatic analyses with assistance from K.A.R. N.W. assisted with neuropathology concerns. N.K.F., R.F. T.A.R, R.R.G and A.M.D. were responsible for the design of the study. All authors assisted with manuscript preparation.

