## Supplementary Figures for "Spatial transcriptomic analysis of childhood ependymoma implicates unresolved wound healing as a driver of tumor progression"

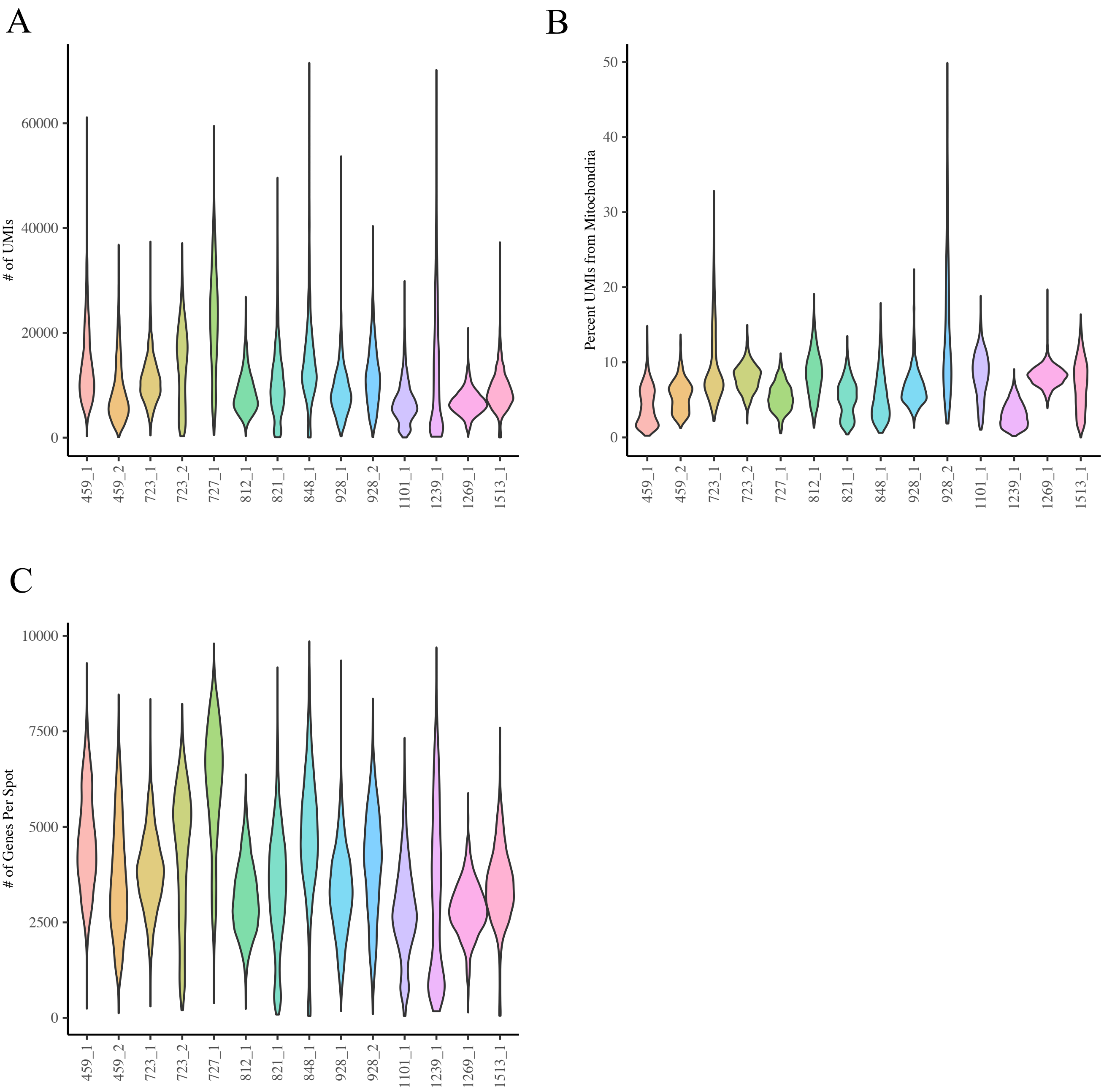


**Supplementary figure 1. Quality control of Visium spatial transcriptomic data.** **A**, Post-filtering number of unique molecular identifiers (UMI) per cell and sample. **B**, percent mitochondrial reads per cell and sample. **C**, number of genes per cell per sample

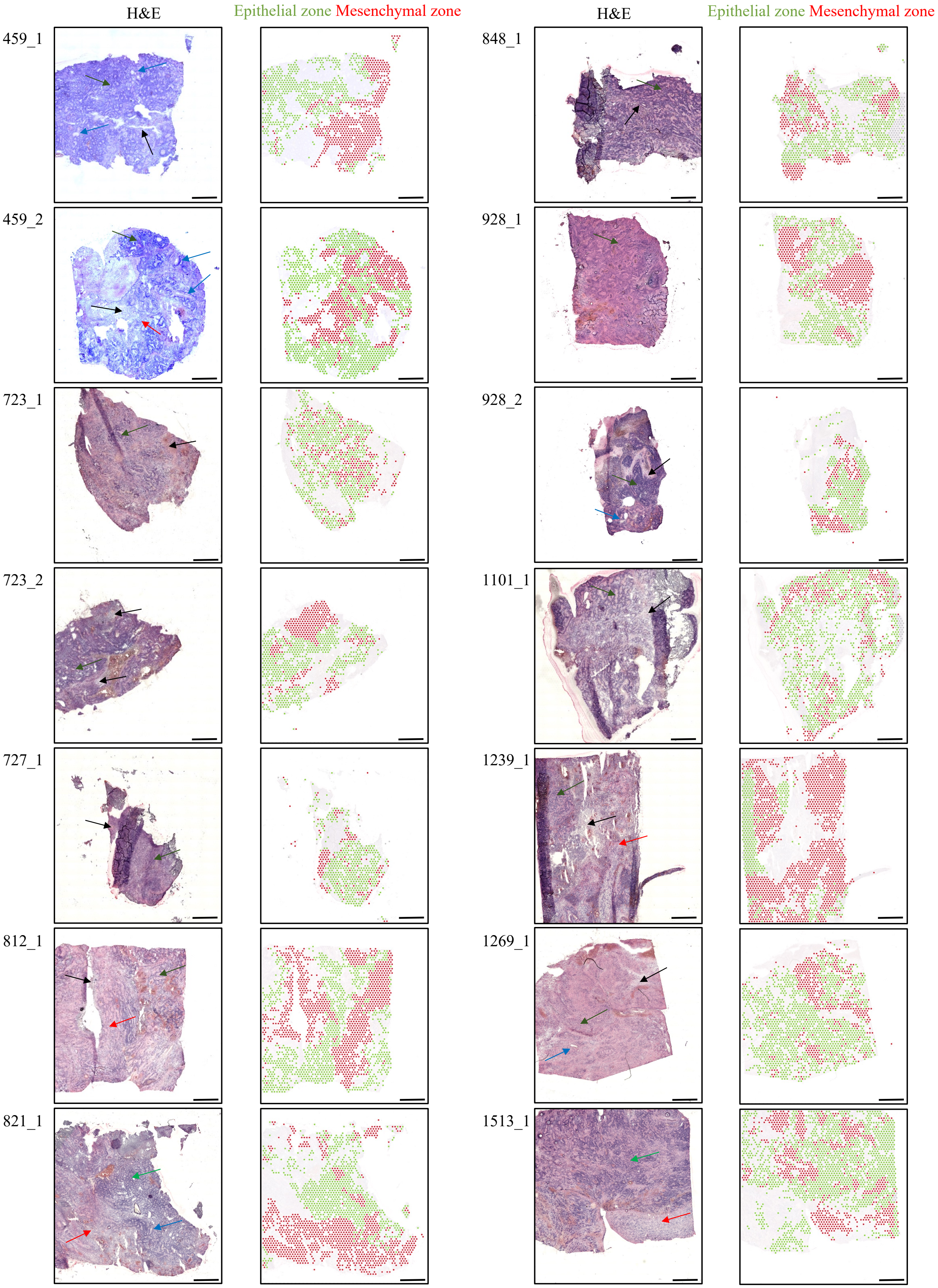

H&E
Epithelial zone Mesenchymal zone
H&E
Epithelial zone Mesenchymal zone
459_1
459_2
723_1
723_2
727_1
812_1
821_1
848_1
928_1
928_2
1101_1
1239_1
1269_1
1513_1

**Supplementary figure 2. Mesenchymal and transcriptomic zone in PFA tumor microenvironment.** Histology (H&E) (left panel) and overlayed epithelial and mesenchymal zone clusters across the tumor microenvironment (right panel) in 14 PFA samples. Histology of PFAs include regions of hypercellularity with prominent perivascular pseudorosettes (blue arrows), areas of increased true ependymal rosettes and epithelial differentiation (green arrows), paucicellular regions with a lesser degree of differentiation (red arrow), and areas of necrosis (black arrow). Size bar = 1mm.

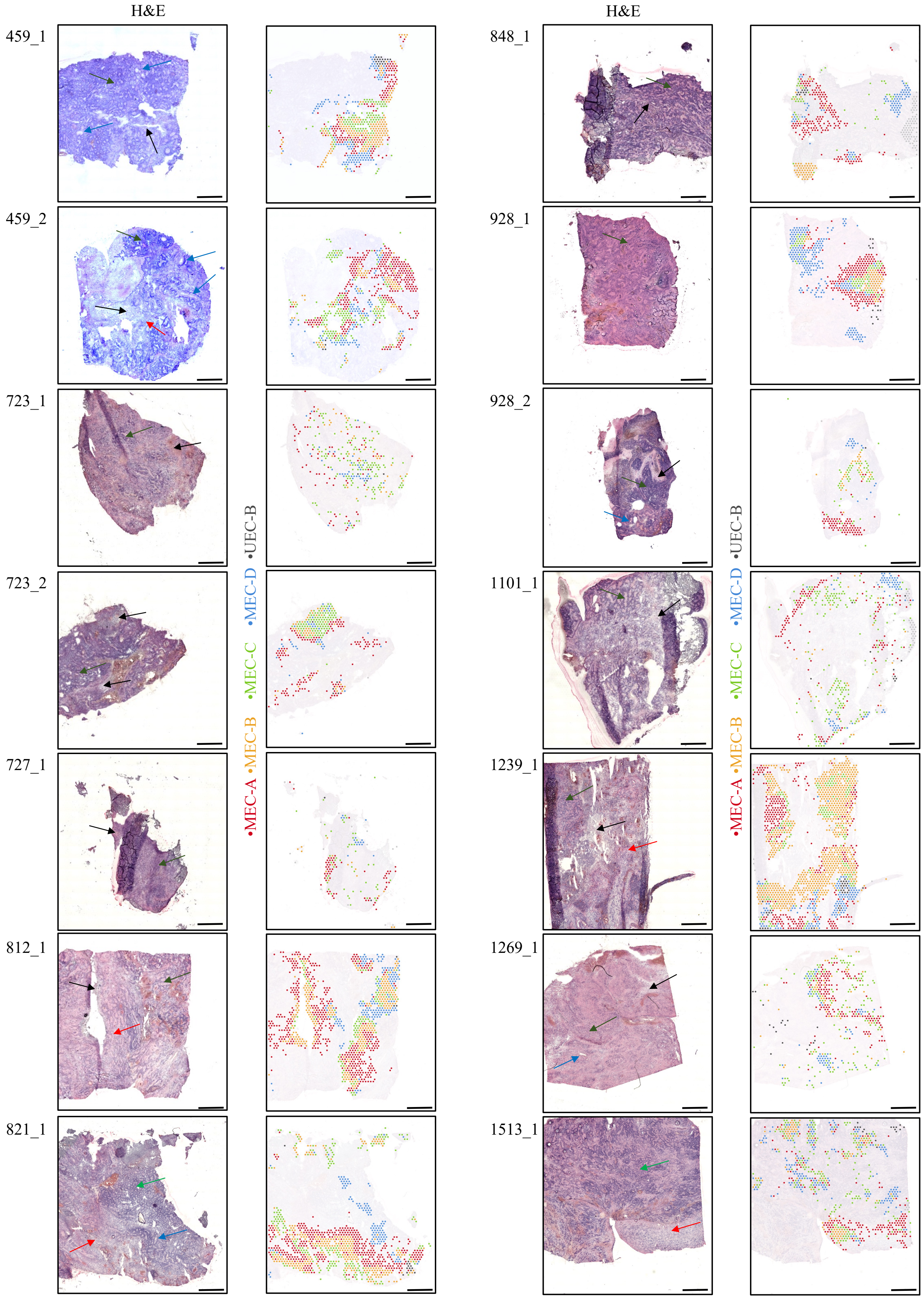
H&E
H&E
459_1
459_2
723_1
723_2
727_1
812_1
821_1
848_1
928_1
928_2
1101_1
1239_1
1269_1
1513_1
•MEC-A •MEC-B •MEC-C •MEC-D •UEC-B
•MEC-A •MEC-B •MEC-C •MEC-D •UEC-B

**Supplementary figure 3. Spatial arrangement of mesenchymal zone subclusters in PFA.** Histology (H&E) (left panel) and overlayed mesenchymal zone subclusters across the tumor microenvironment (right panel) in 14 PFA samples. Histology of PFAs include regions of hypercellularity with prominent perivascular pseudorosettes (blue arrows), areas of increased true ependymal rosettes and epithelial differentiation (green arrows), paucicellular regions with a lesser degree of differentiation (red arrow), and areas of necrosis (black arrow). Size bar = 1mm.

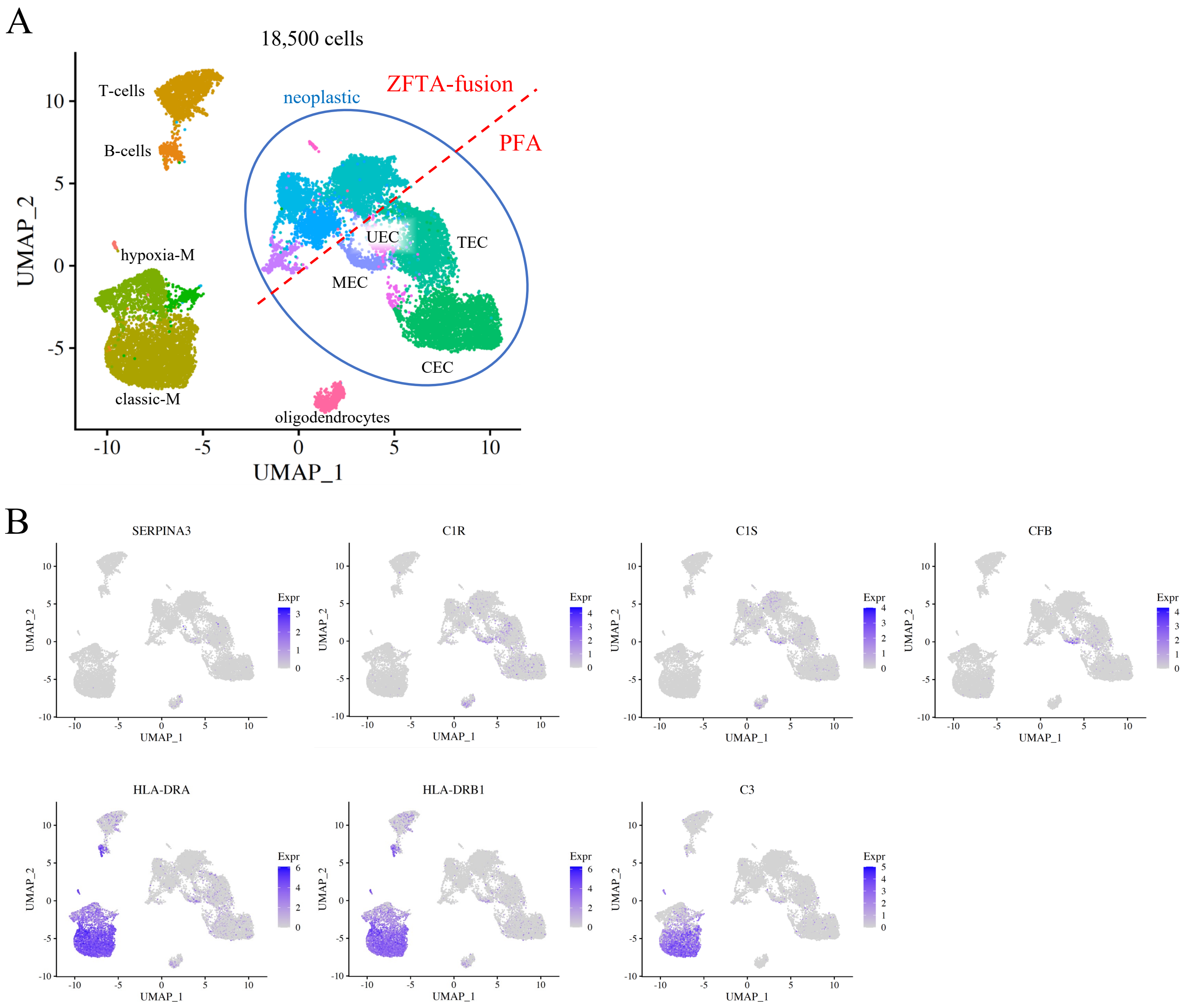


**Supplementary Figure 4. MEC-A marker gene distribution in ependymoma scRNAseq data. A**, Reference UMAP projection of scRNAseq data from 26 EPN patient samples with major cell types and subgroups indicated. **B**, UMAP feature plots of top MEC-A Visium cluster marker expression in scRNAseq dataset.

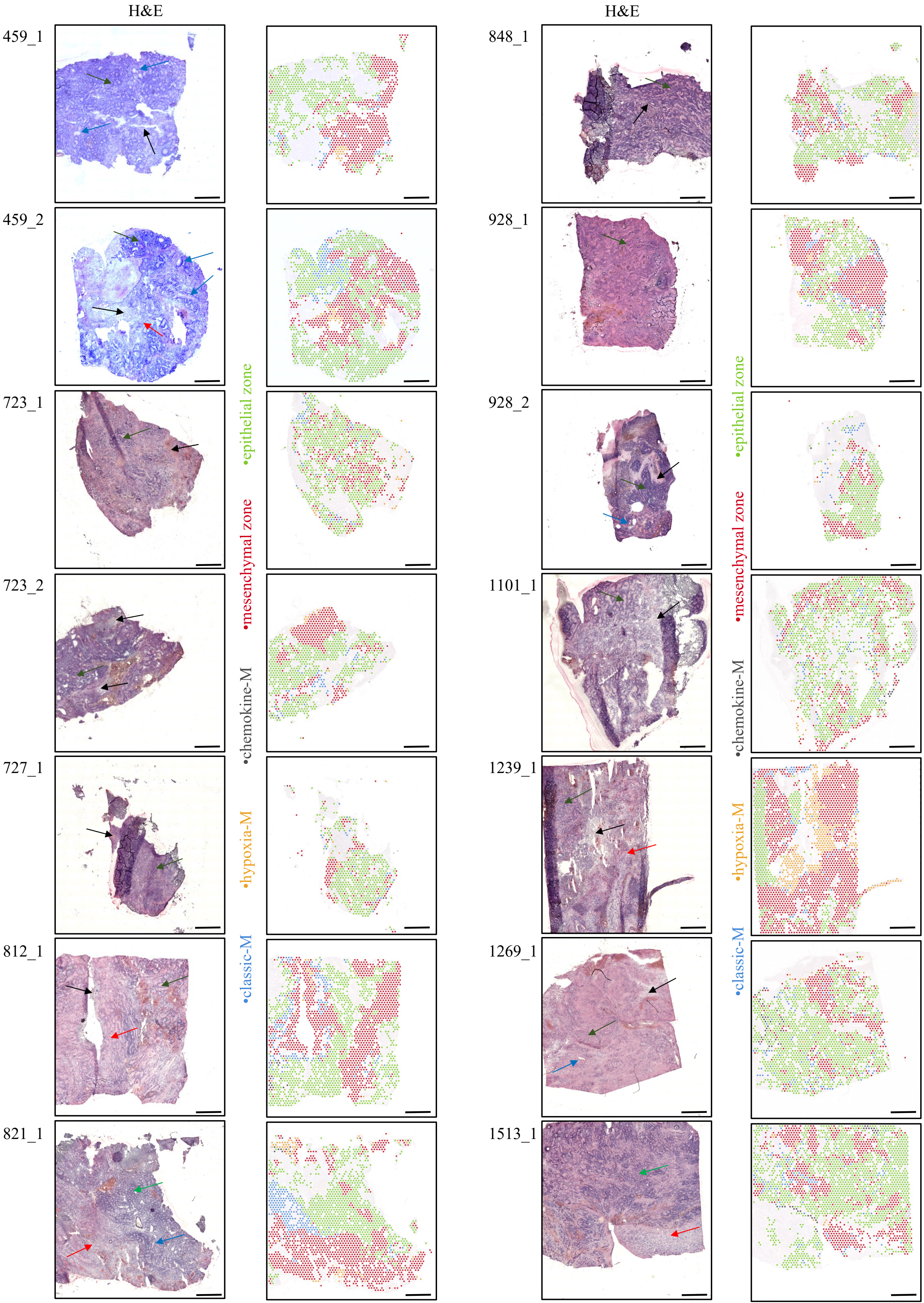
H&E
H&E
459_1
459_2
723_1
723_2
727_1
812_1
821_1
848_1
928_1
928_2
1101_1
1239_1
1269_1
1513_1
•epithelial zone
•mesenchymal zone
•chemokine-M
•hypoxia-M
•classic-M

**Supplementary Figure 5. Spatial arrangement of myeloid cell clusters in relation to epithelial and mesenchymal zone clusters in PFA.** Histology (H&E) (left panel) and overlayed immune, epithelial and mesenchymal zone clusters across the tumor microenvironment (right panel) in 14 PFA samples. Histology of PFAs include regions of hypercellularity with prominent perivascular pseudorosettes (blue arrows), areas of increased true ependymal rosettes and epithelial differentiation (green arrows), paucicellular regions with a lesser degree of differentiation (red arrow), and areas of necrosis (black arrow). Size bar = 1mm.

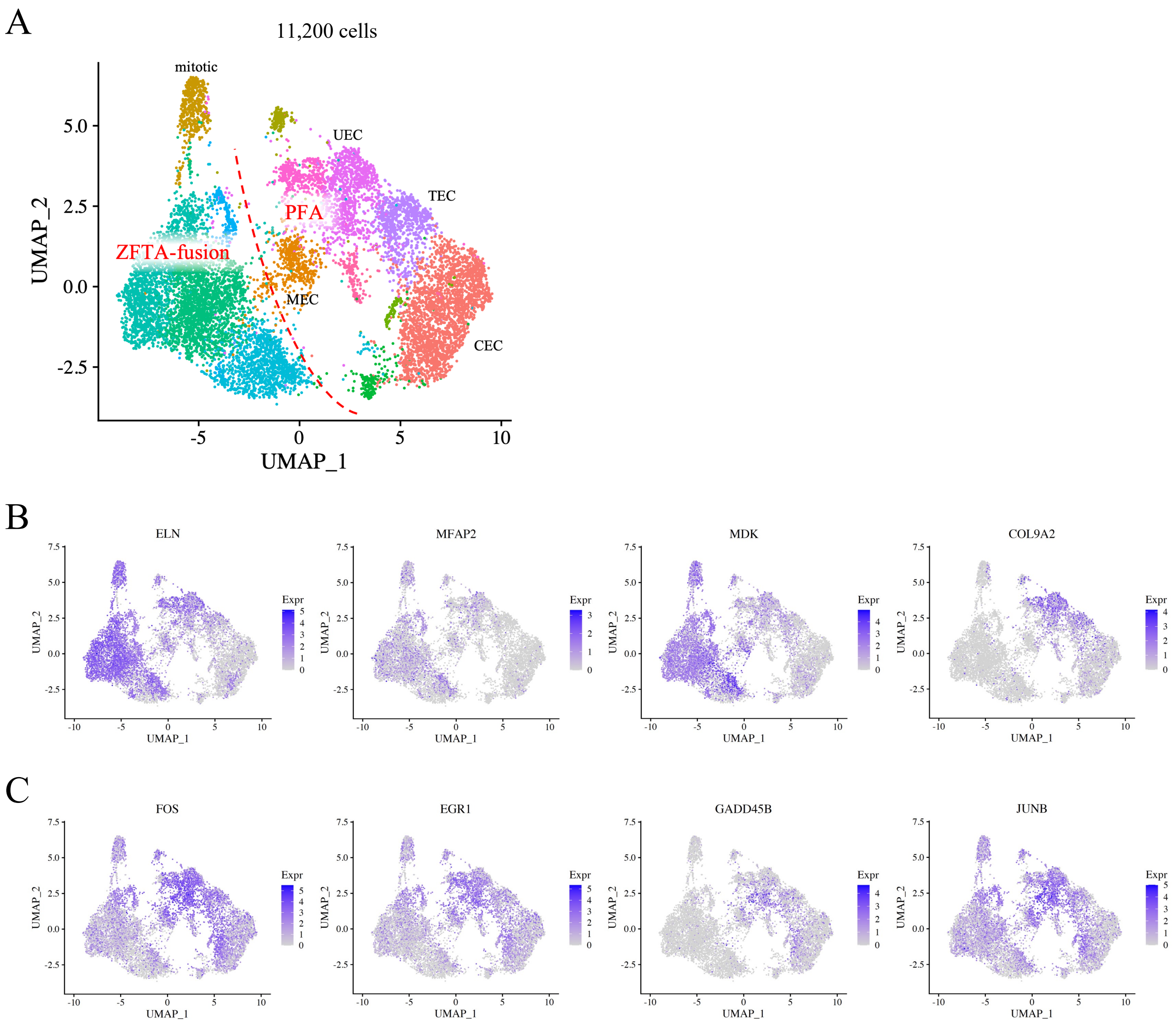


**Supplementary figure 6. ST cluster UEC-A and UEC-B marker gene distribution in ependymoma scRNAseq data. A**, Reference UMAP projection of neoplastic scRNAseq dataset from 26 EPN patient samples with subgroup and major PFA cell types indicated. UMAP feature plots of top **B**, UEC-A and **C**, UEC-B ST cluster marker expression in scRNAseq dataset.

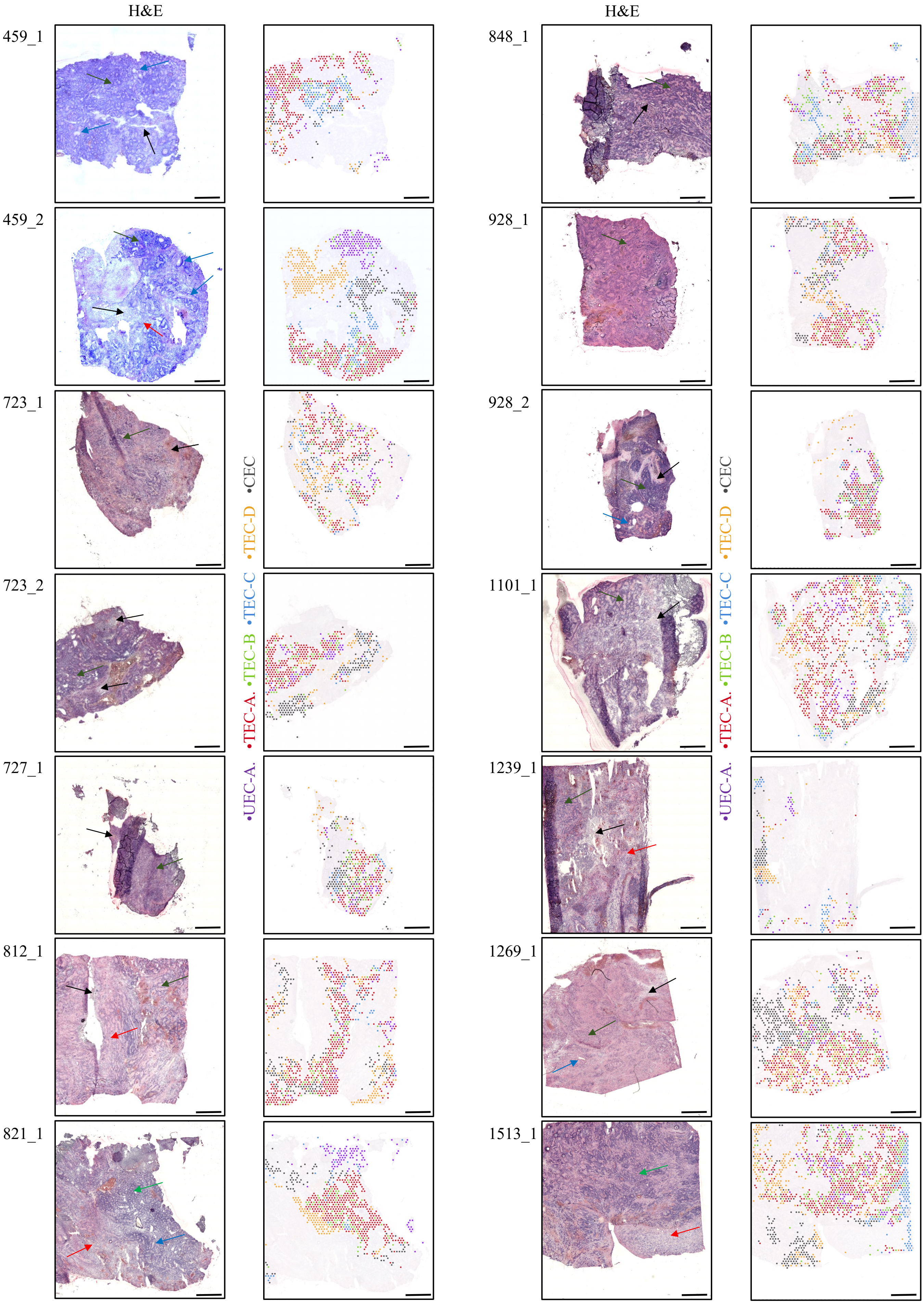
H&E
H&E
459_1
459_2
723_1
723_2
727_1
812_1
821_1
848_1
928_1
928_2
1101_1
1239_1
1269_1
1513_1
•UEC-A. •TEC-A. •TEC-B •TEC-C •TEC-D •CEC
•UEC-A. •TEC-A. •TEC-B •TEC-C •TEC-D •CEC

**Supplementary Figure 7. Spatial arrangement of epithelial zone subclusters in PFA.** Histology (H&E) (left panel) and overlayed epithelial zone subclusters across the tumor microenvironment (right panel) in 14 PFA samples. Histology of PFAs include regions of hypercellularity with prominent perivascular pseudorosettes (blue arrows), areas of increased true ependymal rosettes and epithelial differentiation (green arrows), paucicellular regions with a lesser degree of differentiation (red arrow), and areas of necrosis (black arrow). Size bar = 1mm.
